# Engineering Viral Vectors for Acoustically Targeted Gene Delivery

**DOI:** 10.1101/2021.07.26.453904

**Authors:** Hongyi Li, John E. Heath, James S. Trippett, Mikhail G. Shapiro, Jerzy O. Szablowski

**Author notes:** Correspondence should be addressed to M.G.S. and J.O.S.

## Abstract

Targeted gene delivery to the brain is a critical tool for neuroscience research and has significant potential to treat human disease. However, the site-specific delivery of common gene vectors such as adeno-associated viruses (AAVs) is typically performed via invasive injections, limiting their scope of research and clinical applications. Alternatively, focused ultrasound blood-brain-barrier opening (FUS-BBBO), performed noninvasively, enables the site-specific entry of AAVs into the brain from systemic circulation. However, when used in conjunction with natural AAV serotypes, this approach has limited transduction efficiency, requires ultrasound parameters close to tissue damage limits, and results in undesirable transduction of peripheral organs. Here, we use high throughput *in vivo* selection to engineer new AAV vectors specifically designed for local neuronal transduction at the site of FUS-BBBO. The resulting vectors substantially enhance ultrasound-targeted gene delivery and neuronal tropism while reducing peripheral transduction, providing a more than ten-fold improvement in targeting specificity. In addition to enhancing the only known approach to noninvasively target gene delivery to specific brain regions, these results establish the ability of AAV vectors to be evolved for specific physical delivery mechanisms.

## INTRODUCTION

Gene therapy is one of the most promising emerging approaches to treating human disease. Recently, a number of gene therapies were approved for clinical use, including blindness^1^, muscular dystrophy^2^, and metabolic disorders^3^. Many of these therapies use adeno-associated viral vectors (AAVs) to deliver genes to various organs, but few target the brain. Although several neurological and psychiatric diseases could benefit from gene therapies targeting specific neural circuits, a key challenge limiting the development of such treatments is the need for invasive intracranial injections of the viral vectors. While recent advances are enabling brain-wide gene delivery from systemic^4–6^ or cerebrospinal fluid circulation^7^, these approaches do not provide the spatial targeting needed to address regionally defined neural circuits.

Focused ultrasound blood-brain barrier opening (FUS-BBBO) is a recently developed technique with the potential to overcome these limitations by providing a route to noninvasive, site-specific gene delivery to the brain^8–12^. In FUS-BBBO ultrasound is focused through an intact skull^13, 14^ to transiently loosen tight junctions in the BBB and allow for the passage of AAVs from the blood into the targeted brain site. FUS-BBBO can target intravenously administered AAVs to millimeter-sized brain sites or cover large regions of the brain without tissue damage. These capabilities place FUS-BBBO in contrast with intracerebral injections, which are invasive and deliver genes to a single 2-3 millimeter-sized region per injection^15, 16^, requiring a large number of brain penetrations to cover larger regions of interest. At the same time, the spatial targeting capability of FUS-BBBO differentiates it from the use of spontaneously brain-penetrating engineered AAV serotypes, which lack spatial specificity^5^. In proof of concept studies, FUS-BBBO has been used in rodents to introduce AAVs encoding reporter genes such as GFP^8, 9^, growth factors^17^, optogenetic receptors^10^. The delivery of chemogenetic receptors to the hippocampus provided the ability to modulate memory formation^11^. Despite its promise, three critical drawbacks currently limit the potential of FUS-BBBO in research and therapy applications. First, while the BBB effectively prevents non-FUS-targeted regions of the brain from transduction by systemically administered AAV, peripheral organs have endothelia that allow AAV entry and consequently receive a high dose of the virus, which could lead to toxicity^18^. Second, the relative inefficiency of AAV entry at the site of FUS-BBBO leads to the requirement of high doses of systemic AAVs, on the order of ^~^10^10^ viral particles per gram of body weight. While this magnitude has been used in recent clinical trials, it drives higher peripheral transduction and adds to the cost of potential therapies. Third, efficient delivery of AAV typically require acoustic parameters below, but close to^9, 19^, the threshold for brain tissue damage, reducing the margin for error in interventional planning. We reasoned that these limitations arise from the fact that wild-type serotypes of AAV did not evolve to cross physically loosened biophysical barriers and are therefore not optimal for this purpose. We hypothesized that we could address these limitations by developing new engineered viral serotypes specifically optimized for FUS-BBBO delivery. Capsid engineering techniques^20^ in which mutations are introduced into viral capsid proteins have been used to enhance gene delivery properties such as tissue specificity^5, 6, 21–23^, immune evasion^24^, or axonal tracing^25^. However, they have not yet been used to optimize viral vectors to work in conjunction with specific physical delivery mechanisms.

To test our hypothesis, we performed in vivo selection of mutagenized AAVs in mice in conjunction with FUS-BBBO (**Fig. 1**) by adapting a recently developed Cre-recombinase-based screening methodology^6, 23^. We identified 5 viral capsid mutants with enhanced transduction at the site of FUS-BBBO and but not in the untargeted brain regions. We then performed detailed validation experiments comparing each of these mutants to the parent wild-type AAV, revealing a significant increase in on-target transduction efficiency, increased neuronal tropism, and a marked decrease in off-target transduction in peripheral organs, resulting in an overall performance improvement of more than 10-fold. These results demonstrate the evolvability of AAVs for specific physical delivery methods.

**Figure 1.**
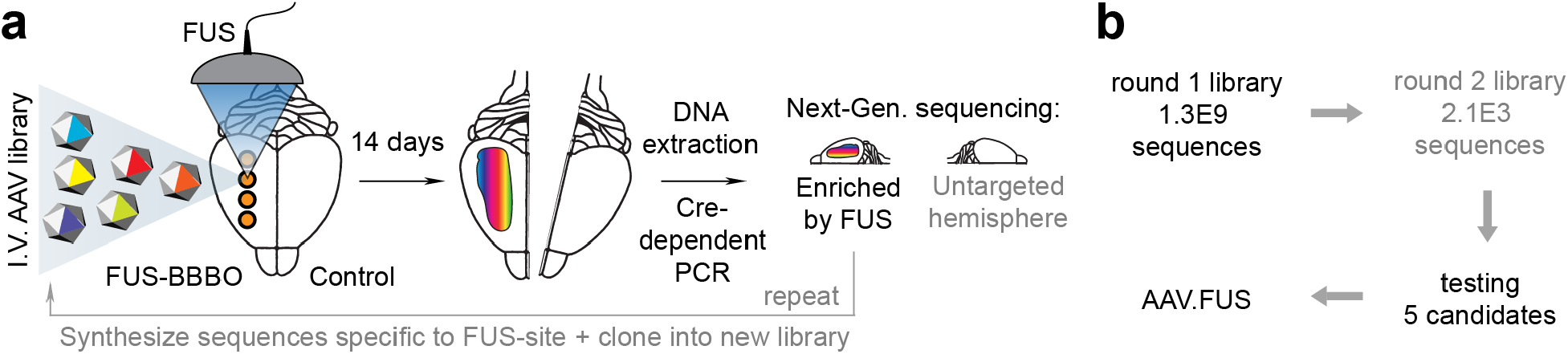
Screening methodology for generation of an AAV for improved site-specific noninvasive gene delivery to the brain. **a**, Summary of the high-throughput screening and selection process. AAV library is administered intravenously (I.V.) and delivered to one brain hemisphere through FUS-BBBO. After 14 days mice are euthanized, their brain harvested, and the DNA from selected hemispheres is extracted. The DNA is then amplified by Cre-dependent PCR that enriches the viral DNA modified by Cre. In our case, neurons expressed Cre exclusively, and the Cre-dependent PCR enriched viral DNA of AAVs that transduced neurons. We subjected the obtained viral DNA to next-generation sequencing for the targeted hemisphere (round 1) or both targeted and control hemispheres (round 2). The process is then repeated for the next round (steps exclusive to round 2 indicated by the grey text). **b**, Overall, 1.3 billion clones were screened in the first round, and 2098 clones in the second round of selection. Out of these clones, we selected 5 that were tested in low-throughput to yield AAV.FUS.3 – a vector with enhanced FUS-BBBO gene delivery.

## Results

### High-throughput in vivo screening for AAVs with efficient FUS-BBBO transduction

To identify new AAV variants with improved FUS-BBBO-targeted transduction of neurons, we generated a library of viral capsid sequences containing insertions of 7 randomized amino acids between residues 588 and 589 of the AAV9 capsid protein (**Supplementary Fig. S1**). AAV9 was chosen as a starting point due to its use in previous FUS-BBBO studies^8, 9, 11^. Meanwhile, 7-mer insertions have been widely used to engineer AAVs with new properties^5, 6, 20–25^.

To make the screening more efficient, we employed recombination-based AAV selection^6, 23^. In this approach, each viral vector genome in the library encodes a mutated capsid protein alongside another segment of DNA that gets inverted in the presence of Cre recombinase (**Supplementary Fig. S1a**). Screening in Cre-expressing cells then allows the identification of capsid variants with the confirmed ability to transduce the cells, with the successful capsid sequences amplified by PCR using inversion-specific primers (**Supplementary Fig S1b**). Using this approach, *in vivo* selection in mice expressing Cre recombinase under neuron-specific promoters ensures the recovery of neuron-transducing AAV variants^5, 6, 26^.

To screen specifically for mutants with enhanced transduction at the site of FUS-BBBO, we followed a two-step strategy (**Fig. 1a, b**). We first down-selected our 1.3 × 10^9^-capsid library to a smaller number of variants that were capable of entering the brain at the site of FUS-BBBO, and then re-screened these variants to quantify enhancement and FUS-target specificity of FUS-BBBO-mediated transduction. We employed FUS parameters below tissue damage limits^11, 27^ (0.33 MPa at 1.5 MHz, 10 ms pulse length, 1 Hz repetition frequency, 0.22 ml dose of microbubbles per gram of body weight). We injected an AAV library intravenously (IV) and immediately after applied FUS to the brains of two hSyn1-CRE mice to open the BBB, using magnetic resonance imaging (MRI) guidance for anatomical targeting, and confirming the opening with contrast-enhanced MRI. We targeted 4 sites within one hemisphere (Fig. 2a). Our library contained a total of 1.3 × 10^9^ clones, at a dose of 6.7×10^9^ viral particles per gram of body weight. After 2 weeks we euthanized the mice and collected their brains. Immediately after, we extracted the viral DNA from the brain and used Cre-dependent PCR amplification to recover and selectively amplify the viral DNA from AAV particles that transduced neurons. We then performed next-generation sequencing (NGS) of the capsid region and selected the 2,098 most abundant sequences for subsequent evaluation. Based on the design of this screen, each of these variants was pre-selected to able to enter the brain at the site of FUS-BBBO and transduce neurons. However, the specificity of FUS-mediated entry and efficiency of transduction by these variants could not be determined quantitatively at this stage due to the small copy number of each variant in the library.

**Figure 2.**
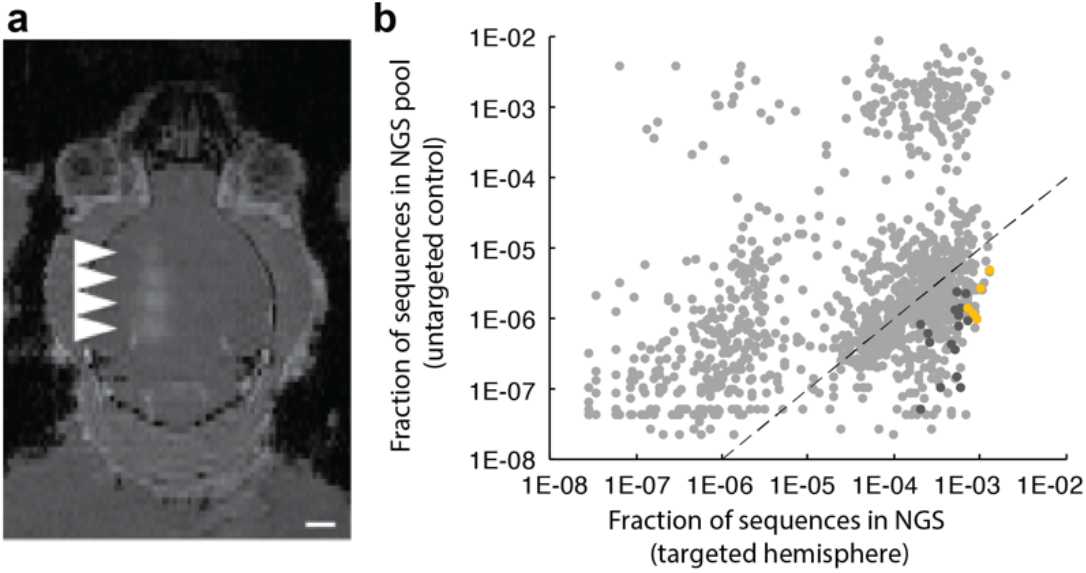
High throughput screening yields vectors with improved FUS-BBBO gene delivery. **a**, An MRI image showing mouse brain with 4 sites opened with FUS-BBBO in one hemisphere. The bright areas (arrowheads) indicate successful BBB opening and extravasation of the MRI contrast agent Prohance into the brain. This BBB opening was used for delivery of the AAV library. **b**, Sequencing results of round 2 of screening show a fraction of NGS reads within the DNA extracted from brains of Syn1-Cre mice subjected to FUS-BBBO and injected with a focused library of 2098 clones. Each dot represents a unique capsid protein sequence, and the position on each axis corresponds to the number of times the sequence was detected in the FUS-targeted and untargeted hemispheres. Markers below the dotted line represent sequences that on average showed 100-fold higher enrichment in the targeted hemisphere as compared to the control hemisphere. Dark grey dots represent 35 clones that are enriched in the FUS targeted hemispheres at least 100-fold in every tested mouse and DNA sequence encoding the 7-mer insertion peptide. Yellow dots represent 5 clones (AAV.FUS.1-5) selected for low-throughput testing. Due to the use of a logarithmic plot, clones that had zero copies detected in either of the hemispheres are not shown.

To quantitatively compare our 2,098 down-selected capsid variants, we re-synthesized and packaged them as a new AAV library at a dose of 1.3 × 10^9^ viral particles per gram of body weight, corresponding to ^~^1.5-3×10^7^ viral particles of each clone being injected into each mouse. In each of two hSyn-CRE mice, we injected an AAV library intravenously and opened the BBB in one hemisphere using MRI-guided FUS as in round 1. After 2 weeks, the brains were extracted, hemispheres separated and the DNA from FUS-targeted and untargeted hemispheres extracted for each mouse. The DNA extract was amplified by the CRE-dependent PCR to enrich for viral genomes that transduced neurons. After FUS-BBBO delivery, DNA extraction, CRE-dependent PCR, and NGS, we recovered 1,433 sequences.

To identify the most improved candidates we examined their copy number in the targeted hemispheres and compared it to the untargeted hemispheres (**Fig. 2b**). We first looked for variants that were at least 100-fold more represented in the targeted hemisphere relative to the untargeted hemisphere to identify AAVs that selectively transduced sites subjected to FUS-BBBO. From this list, we further selected candidates for which the 100-fold difference was maintained in both mice and in each alternative codon sequence corresponding to its 7-mer peptide, thus ensuring robustness of the NGS data. In the end, 35 sequences met these criteria (dark grey symbols in **Fig. 2b**). Among these FUS-BBBO-specific variants, we chose the 5 most frequently detected sequences, which we hypothesized would code for AAV capsids with the most efficient neuronal transduction. We re-synthesized these sequences, cloned them into the AAV9 capsid between amino acids 587-588, and packaged them for detailed evaluation, naming them AAV.FUS 1 through 5.

### AAV.FUS candidates show enhanced transduction of neurons in targeted brain regions and reduced transduction of peripheral organs

An ideal AAV vector for ultrasound-mediated gene delivery to the brain would efficiently transduce targeted neurons while avoiding peripheral tissues, such as the typically highly transduced liver^28^. Additionally, such a vector should only transduce the brain at the FUS-targeted sites. Of the natural AAV serotypes, AAV9 is most commonly used in FUS-BBBO because it transduces neurons at the ultrasound target with relatively high specificity compared to untargeted brain regions^8, 10, 11^. However, AAV9 requires high doses to achieve significant gene expression and transduces peripheral tissues in the process^8, 10, 11^, leaving substantial room for improvement. We decided to test our five selected AAV.FUS vectors against AAV9.

We performed FUS-BBBO while intravenously co-administering each AAV.FUS candidate alongside AAV9 in individual comparison experiments. Consequently, each mouse had an internal control where the location and extent of FUS-BBBO was identical for both serotypes. To quantify the transduction efficiency, we encoded the fluorescent proteins mCherry and EGFP in AAV9 and each AAV.FUS variant, respectively, under a general CaG promoter. After 2 weeks of expression, we counted the numbers of mCherry and EGFP-expressing cells within the site of FUS-BBBO. The reliability of this quantification method was established by comparing cell counts for co-administered AAV9-EGFP and AAV9-mCherry (**Supplementary Fig. S2**). Our quantification showed that AAV.FUS.1, 2, 3, and 5 had significantly improved transduction efficiency compared to AAV9 (p = 0.0274, 0.0003, 0.0052, 0.0087, respectively, **Fig. 3, a–b**) wherease AAV.FUS.4 showed no improvement (p = 0.2556). The fold-change in transduction relative to AAV9 was greatest for AAV.FUS.2, and lowest for AAV.FUS.4 (**Supplementary Fig. S3**). None of the AAV.FUS candidates produced off-target expression within the brain (**Fig. 3c**).

**Figure 3.**
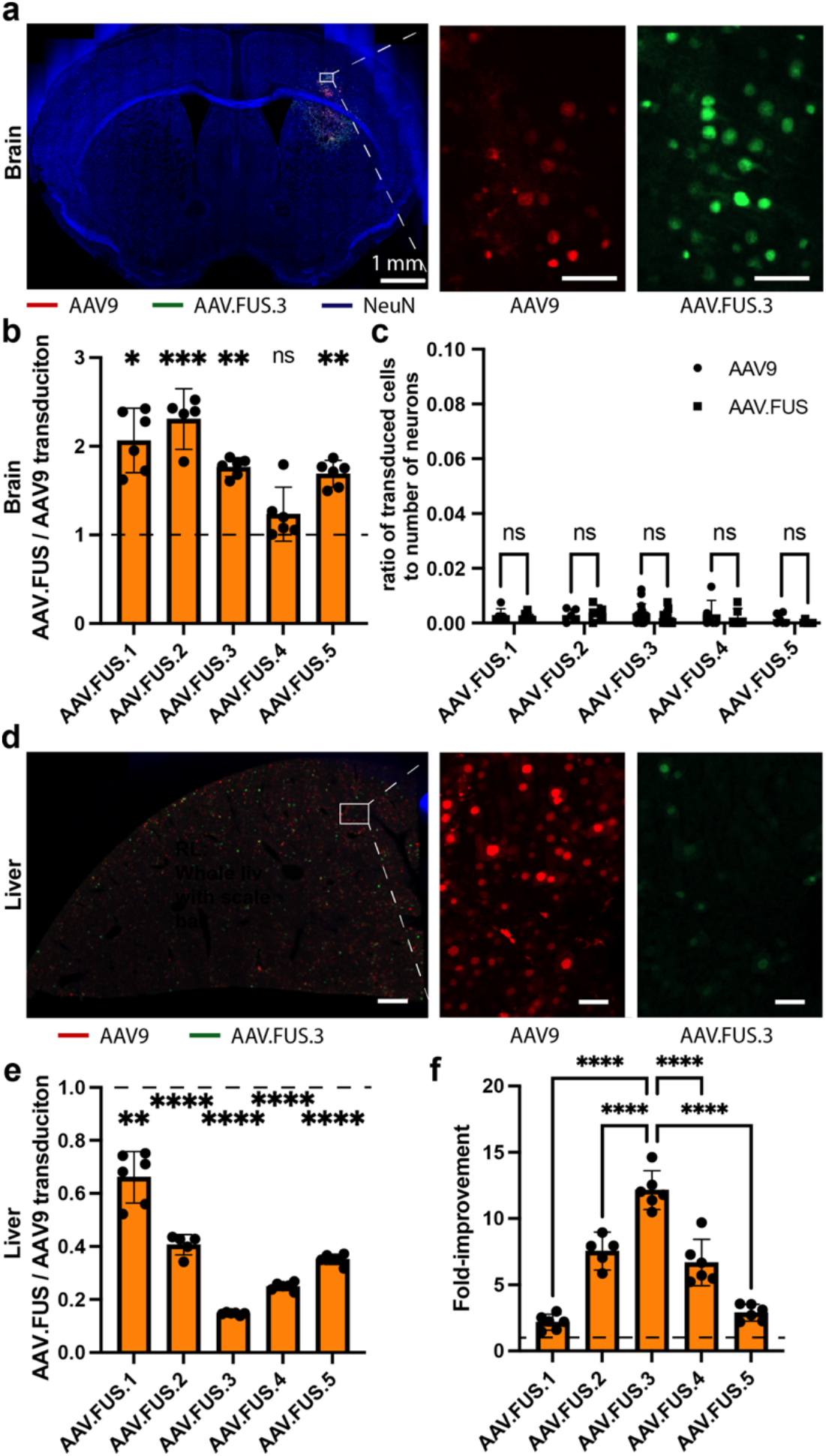
AAV.FUS candidates improve efficiency of gene delivery to the brain and reduce peripheral transduction. **a**, Representative images were obtained from mice co-injected with AAV9 and a AAV.FUS.3 at 1010 viral particles per gram of body weight. After 3 weeks, the mice were perfused, brains were extracted and then sectioned at 50 microns. Sections were imaged on a confocal microscope with 20x objective showing brain transduction by AAV9 (red) and AAV.FUS.3 (green), and counterstained with a neuronal stain (NeuN, blue). **b**, All but one (AAV.FUS.4) AAV.FUS candidates showed significant improvement over the co-injected AAV9. **c**, We found that few cells were transduced outside of the FUS-targeted site and AAV.FUS.3 and AAV9 were not significantly different (0.19% off target transduction for AAV.FUS.3 vs 0.4% for AAV9; p=0.072, two way Anova with Sidak multiple comparisons test). Similarly other candidates also showed no differences (AAV.FUS.1, p=0.99 n=6; AAV.FUS.2, p=0.98, n=5; AAV.FUS.4, p=0.86, n=6; AAV.FUS.5, p=0.83, n=6). **d**, Representative images showing liver transduction by AAV9 (red) and AAV.FUS.3 (green). **e**, All tested candidates showed reduction of the liver transduction as compared to the co-injected AAV9 in the same mice for which brain expression was analyzed. **f**, We defined the fold-improvement in targeting efficiency as the ratio of brain transduction to the liver transduction efficiency using AAV9 as a baseline, which suggested that AAV.FUS.3 is the top candidate for further study. Scale bars are 50 microns in panels a, c, unless otherwise noted. (**** = p<0.0001; *** = p<0.001; ** = p <0.01; * = p<0.05, ns = not significant); Error bars are 95% CI.

Next, we evaluated the extent to which AAV.FUS candidates transduce off-target peripheral organs. In mice that received intravenous co-injections of AAV9-mCherry and each variant of AAV.FUS-EGFP we counted transduced cells in the liver, a peripheral organ known to be targeted by AAVs and a potential source of dose-limiting toxicity^29, 30^. Two weeks after injection, we collected liver tissues and imaged the livers, counting cells expressing each fluorophore (**Fig. 3, d–e**). We found markedly reduced liver transduction among the AAV.FUS candidates compared to AAV9 (**Fig 3 e**). AAV.FUS 3 showed the largest reduction in liver transduction compared to the wild-type serotype (6.8-fold reduction, p<0.0001), which was significantly higher reduction compared to the other tested AAV.FUS candidates (**Supplementary Fig. S4**).

Our analyses of brain and liver transduction showed that AAV.FUS candidates both decrease the targeting of the peripheral tissue and increase the transduction efficiency of the targeted brain regions, which leads to a large overall improvement in transduction specificity, expressed as the ratio of the fold-increase in brain transduction and the fold-decrease in liver transduction compared to AAV9. By this metric, AAV.FUS.3 showed a 12.1-fold improvement, significantly greater than the other candidates (p<0.0001 for all pairwise comparisons, one-way ANOVA with Tukey-HSD post hoc test; **Fig. 3f**).

A final criterion for successful gene delivery in many applications is the ability to transduce specific cell-types at the targeted anatomical location, for example neurons. AAV9 transduces both neuronal and non-neuronal cell types^31–33^. We hypothesized that, since our Cre-dependent screen used mice with the recombinase expressed under a neuronal promoter, our engineered variants could have a higher neuronal tropism relative to their wild-type parent serotype. To test this hypothesis, we immunostained brain sections from mice co-transduced with AAV9-mCherry and each variant of AAV.FUS-EGFP during FUS-BBBO for the neuronal marker NeuN and imaged these sections for GFP, mCherry, and NeuN signal. The fraction of AAV9-transduced (mCherry-positive) cells that were also positive for NeuN was 44.7% (±0.75%, SEM; n=8). In contrast, all AAV.FUS candidates had higher neuronal tropism (**Fig. 4**), with neurons constituting between 64.6 (±0.97%, SEM) (AAV.FUS.1) and 69.8 (±1.8%, SEM) (AAV.FUS.3) of all transduced cells (one-way ANOVA with Tukey-HSD post hoc test; p<0.0001 for all AAV.FUS candidates). These results show that in addition to improved specificity for targeted regions of the brain, the engineered viral capsids are more selective for neurons over other cephalic cell types.

**Figure 4.**
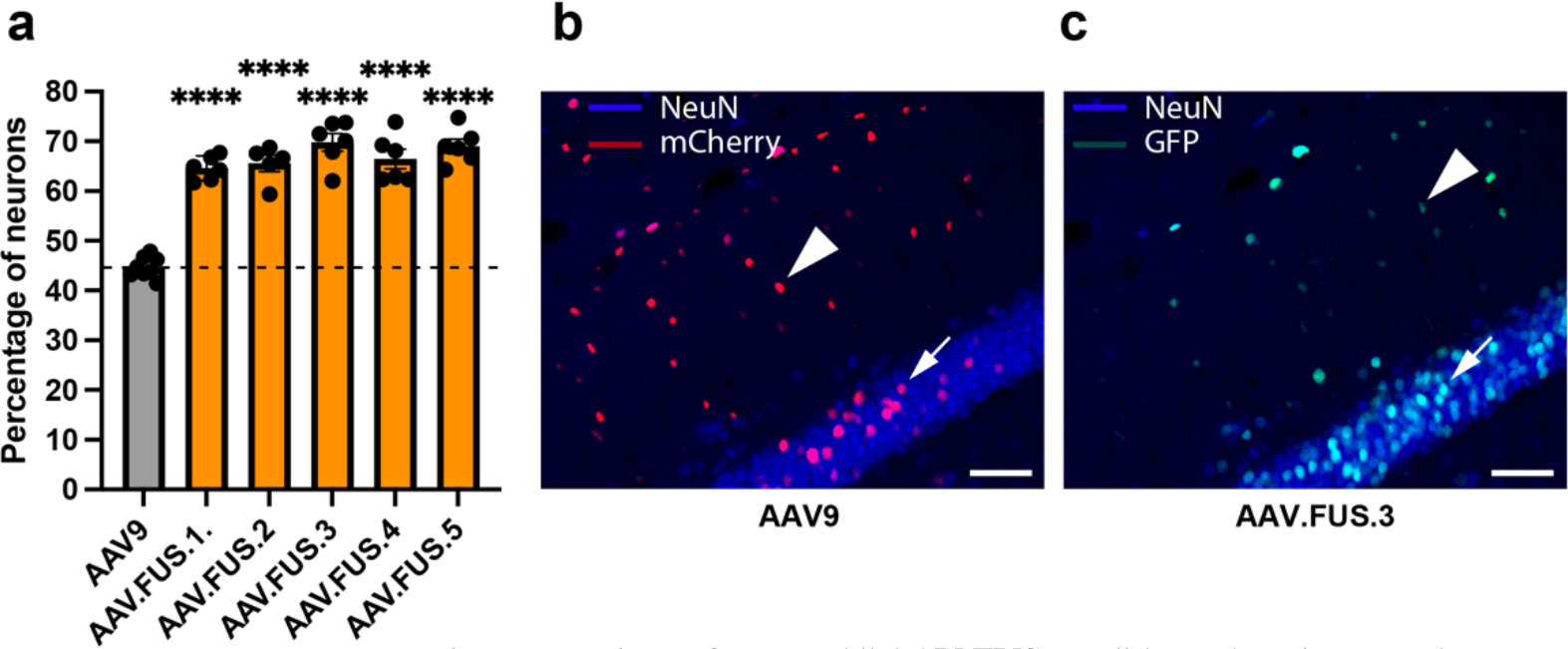
AAV.FUS candidates show improved neuronal tropism. **a**, All AAV.FUS candidates show improved neuronal tropism (p<0.0001). Upon FUS-BBBO gene delivery, AAV.FUS.3. has 56% more likelihood of transducing a neuron than AAV9 (69.8%, vs 44.9% neuronal transduction, respectively; p<0.0001). **b**, Representative images showing AAV9 transducing both neurons (blue, NeuN staining, example neuron designated by an arrow) and non-neuronal cells (example non-neuronal cell designated by an arrowhead), **c**, In comparison, more of the cells transduced with AAV.FUS (green) are neurons (example neuron designated by an arrow), rather than non-neuronal cells (example cell designated by an arrowhead). Scale bars are 50 microns. (All p-values - **** = p<0.0001). Error bars are 95% CI.

### Region-specific transduction efficiency of AAV.FUS.3

Based on its leading combination of neuronal tropism and improvement in brain specificity among the engineered variants, we selected AAV.FUS.3 for further evaluation as a FUS-BBBO-specific viral vector. To further characterize its performance relative to AAV9, we decided to evaluate the efficiency of delivery when these vectors are targeted to different brain regions. To ensure that each region is targeted exclusively, only one brain region was targeted with FUS-BBBO in each tested mouse. To ensure the rigor of this investigation and account for variability in virus titration^34^, we obtained a new batch of both AAV9 and AAV.FUS.3 and tittered them independently. We evaluated the efficiency of transduction when these vectors were targeted by FUS-BBBO to the striatum (caudate putamen), thalamus, hippocampus, and midbrain of AAV.FUS.3.

As in our earlier experiments, we observed a major improvement in AAV.FUS.3 transduction compared to AAV9 in all targeted regions, with a fold-change ranging from 2.4±0.08 to 4.3±0.08 (95% CI, **Fig. 5**). Among brain regions, we found that the hippocampus (Hpc) is transduced with a particularly elevated relative efficiency, while the cortex (Ctx) showed the lowest, but still substantial, improvement. These results indicate that AAV.FUS.3 can target multiple brain regions with improved efficiency, while suggesting the potential for further engineering AAVs with region-enhanced tropism in FUS-BBBO delivery.

**Figure 5.**
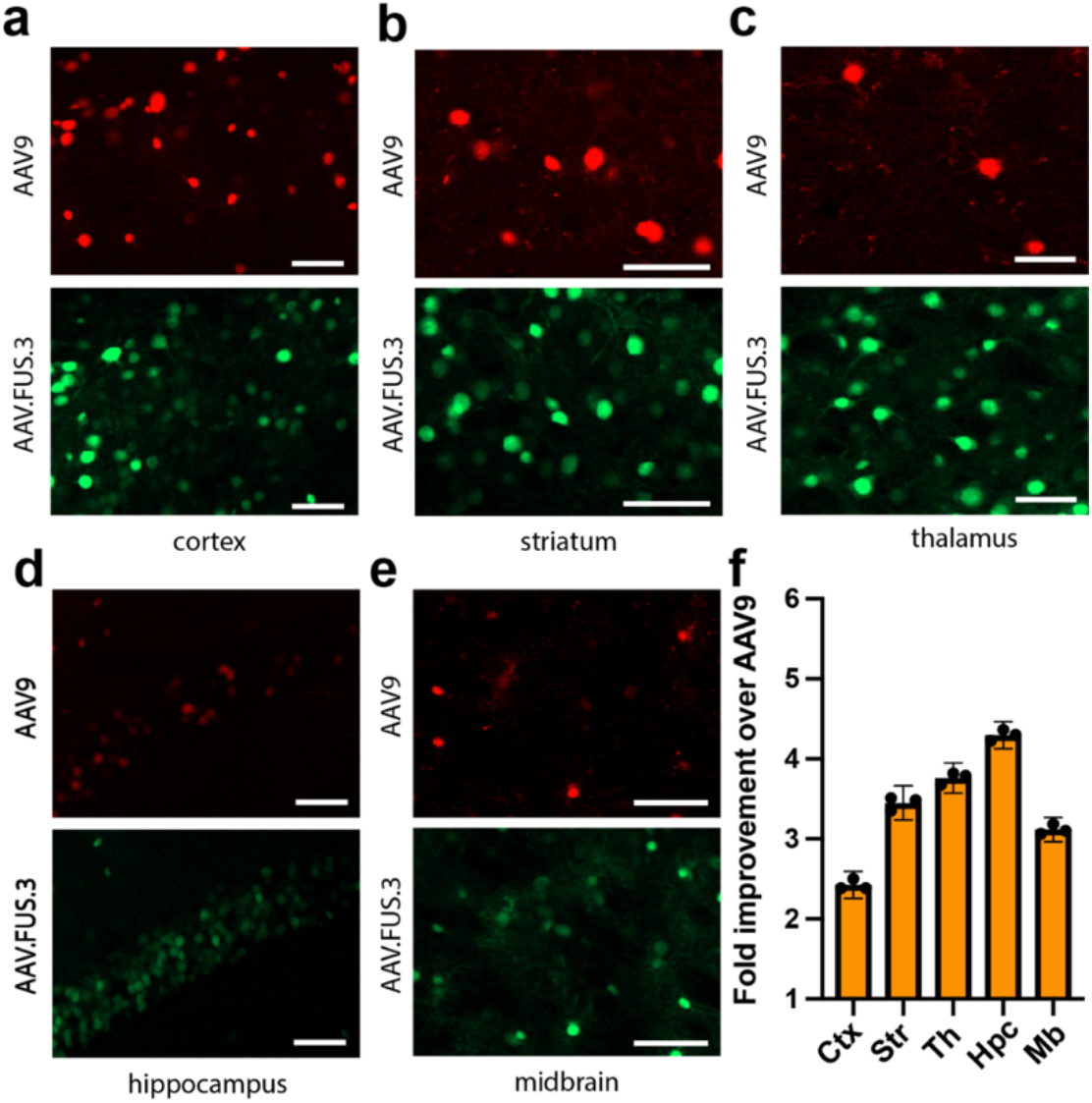
AAV.FUS.3 shows regional dependence of transduction efficiency. Hippocampus showed the highest, 4.3-fold, improvement in transduction over AAV9. **a**, Representative image comparing transduction of the cortex with AAV.FUS.3 (green) and AAV9 (red). **b**, Representative image comparing transduction of the striatum with AAV.FUS.3 (green) and AAV9 (red). **c**, Representative image comparing transduction of the thalamus with AAV.FUS.3 (green) and AAV9 (red). **d**, Representative image comparing transduction of the hippocampus with AAV.FUS.3 (green) and AAV9 (red). **e**, Representative image comparing transduction of the midbrain with AAV.FUS.3 (green) and AAV9 (red). **f**, AAV.FUS.3 shows regional differences in transduction efficiency of the tested regions – cortext (ctx), striatum (Str), thalamus (Th), hippocampus (Hpc), midbrain (Mb). All differences were statistically significant (one way ANOVA, “F (4, 10) = 283.4”, P<0.0001; All pairwise comparison p-values < 0.01, Tukey HSD post-hoc test). Scale bars are 50 microns. Error bars are 95% CI.

## DISCUSSION

Our results show that viral vectors can be engineered to improve noninvasive, site-specific gene delivery to the brain using ultrasound-mediated blood-brain barrier opening. Gene therapy is widely used in research and is becoming a clinical reality. However, most of the available methods for gene delivery to the brain either lack regional specificity or are invasive and challenging to apply to large brain regions^4–7, 15, 16^. While FUS-BBBO promises to overcome these challenges, its use in conjunction with AAVs encounters challenges of parameter safety and peripheral transduction^35^ which, despite longstanding effort, have not been fully solved through the optimization of ultrasound parameters^8, 36, 37^ and equipment^38, 39^ alone. Simply increasing the intravenous dose of natural AAVs is not feasible due to the additional cost of the virus production, stronger immune response to the virus^40^, higher non-specific transduction^41–43^ of peripheral tissues and associated toxicity^30, 44, 45^.

In this study, we approached the problem of improving FUS-BBBO gene delivery by engineering the viral vectors themselves. The resulting improvements include an increase in brain transduction per virus injected, a reduction in peripheral expression and an increase in neuronal tropism. Among the selected 5 AAV.FUS candidates, four transduced target brain sites more efficiently than AAV9 while also lowering transgene expression in the liver in the same mice. Our top candidate, which we call AAV.FUS.3, demonstrated improved transduction in five different brain regions and an overall efficiency of targeting the brain, defined as the ratio of brain to liver (peripheral) transduction, improved 12.1-fold compared to AAV9. This improvement in tissue specificity is particularly important because peripheral transduction can lead to toxicity. For example, AAV-based gene therapy has been shown to induce liver toxicity in clinical trials^29^.

Our results suggest the need to investigate the mechanisms by which AAVs enter the brain after FUS-BBBO and what accounts for the differences in efficiency among serotypes. The prevailing understanding of FUS-BBBO mechanisms suggests that FUS loosens tight junctions in the vasculature, allowing molecules and nanoparticles such as AAVs to pass from the blood into the brain^46^. Within this framework, reductions in peripheral uptake (leaving more AAV to circulate) and reduced binding to extracellular matrix^47^ could help certain serotypes enter through physically-generated openings and reach neurons more efficiently. Another possibility is that FUS-BBBO could cause molecular changes to the vascular endothelium, leading to a more complex interaction between viral vectors and their target. Understanding these factors would enable additional future engineering and optimization of FUS-BBBO-based gene delivery.

Overall, this study shows that the molecular engineering of AAV capsids can lead to improved ultrasound-mediated gene delivery to the brain. Our screen yielded AAV.FUS.3, the first viral vector expressly engineered to work in conjunction with a specific physical delivery method.

## MATERIALS AND METHODS

### Animals

Animals. C57BL/6J and Syn-1-Cre mice were obtained from Jackson Lab. Animals were housed in a 12h light/dark cycle and were provided with water and food ad libitum. All experiments were conducted under a protocol approved by the Institutional Animal Care and Use Committee (IACUC) of the California Institute of Technology.

### Focused ultrasound equipment and BBB opening procedures

FUS-BBBO. Adult male Syn1-Cre mice (at least 14 weeks old) were anesthetized with 2% isoflurane in air, the hair on their head removed with Nair depilation cream and then cannulated in the tail vein using a 30-gauge needle connected to PE10 tubing. The cannula was then flushed with 10 units (U)/ml of heparin in sterile saline (0.9% NaCl) and attached to the mouse tail using tissue glue (Gluture). Subsequently, the mice were placed in the custom-made plastic head mount and imaged in a 7T MRI (Bruker Biospec). A fast low-angle shot sequence (echo time TE=3.9ms, repetition time TR=15ms, flip angle 20°) was used to record the position of the ultrasound transducer in relation to the mouse brain. Subsequently, the mice were injected via tail vein with AAVs. Immediately after viral injection, the mice were also injected with 1.5×10^6^ DEFINITY microbubbles (Lantheus) and 0.125 μmol of ProHance (Bracco Imaging) dissolved in sterile saline, per g of body weight. The dose of DEFINITY was identical as used in our previous studies^1^. The dose of ProHance was chosen based on the manufacturer’s recommendations. Within 30 s, the mice were insonated using an eight-channel FUS system (Image Guided Therapy) driving an eight-element annular array transducer with a diameter of 25 mm and a natural focal point of 20 mm, coupled to the head via Aquasonic ultrasound gel. The gel was placed on the top and both sides of the animal’s head to minimize reverberations from tissue/air interfaces. The focal distance was adjusted electronically. The ultrasound parameters used were 1.5MHz, 1% duty cycle, and 1Hz pulse repetition frequency for 120 pulses and were derived from a published protocol. The pressure was calibrated using a fiber optic hydrophone (Precision Acoustics), with 21 measurements and uncertainty of ±3.8% (SEM). The pressure for FUS-BBBO was chosen to maximize the safety of delivery and was chosen on the basis of our previous studies^1^ and preliminary data in our laboratory. The ultrasound parameters were 1.5MHz, 0.33 MPa pressure accounting for skull attenuation (18%)^48^, 1% duty cycle, and 1Hz pulse repetition frequency for 120 pulses. For each FUS site, DEFINITY and Prohance were re-injected before the additional insonation. Each animal underwent four insonations located in one hemisphere, starting from the midbrain and going forward. The time between each insonation was approximately 3 minutes and included 120 s of insonation and 1 minute for readjustment of positioning on the stereotaxic frame. The center focus of beams was separated by 1.35 – 1.5 mm (depending on mouse weight 25-35 g) in the anterior / posterior direction.

### Plasmids and DNA library generation

The plasmids used were either obtained from Addgene, Caltech’s vector core, or modified from these plasmids. The AAV library genome used for selection (acceptor plasmid, rAAV9Rx/a-delta-CAP) was obtained from Caltech’s vector core facility, as were other plasmids (REP2-CAP9Stop-DeltaX/A, pUC18). The Rep-Cap plasmid for packaging AAV.FUS candidates were modified from Addgene plasmid #103005 by introducing mutations selected from the screen. For testing the transduction we used a plasmid obtained from Addgene (pAAV-CaG-NLS-EGFP - #104061) and a plasmid modified in-house with exchanged EGFP for mCherry protein (pAAV-CaG-NLS-mCherry).

Mutations were introduced into the acceptor plasmid using a PCR with degenerated primers (7MNN) with a sequence 5’-GTATTCCTTGGTTTTGAACCCAACCGGTCTGCGCCTG TGCNMNNMN NMNNMNNMNNMNNTTGGGCACTCTGGTGGTTTGTG-3’, targeted as a 7-aminoacid insertion between residues 587 and 588. The amplified insert was then introduced into the capsid plasmid through restriction cloning using XbaI and AgeI enzymes. DNA from the treated brain was recovered by PCR using two pairs of plasmids – the first step of amplification was done using 5’-CAGGTCTTCACGGACTCAGACTATCAG-3’ and 5’-CAAGTAAAACCTCTACAAATGTGGTAAAATCG-3’ primers which selected for the DNA that has been modified by Cre enzyme. The second stage, intended to amplify the DNA was performed using a pair of primers: 5’-ACTCATCGACCAATACTTGTACTATCTCTCTAGAAC-3’ and 5’-GGAAGTATTCCTTGGTTTTGAACCCAA-3’.

### Virus production and purification

AAV library was purified as previously published^6^. In short, we transfected the DNA carrying a genome containing capsid which has been modified by the 7-mer insertion (10 ng per 100 mm diameter dish), the helper DNA containing REP protein (10 mg per 100 mm diameter dish, and 9.99 mg of empty pUC19 carrier plasmid), and an AdV helper plasmid (20 mg per 100 mm diameter dish) using PEI. Media was changed 16 h after transfection, and then collected 48 h post-transfection and stored in 4C. 60h after the transfection, we scraped the cells into San digestion buffer (Tris pH 8.5 with 500mM NaCl and 40 mM MgCl2 with Salt Active Nuclease). Virus in the media was precipitated using 1/5 volume of 5X PEG8000+NaCl (40% PEG-8000 and 2.5M NaCl), incubated on ice for 2 h, and spun at 3000xg for 30min at 4C. The media and cell-scraped stocks were then combined and precipitated using iodixanol gradient precipitation (virus appears on the 40-60% iodixanol interface), diluted into 15 ml PBS with 0.001% Pluronic-F68, and sterile-filtered through a 0.2-micron PES filter. Finally, the buffer was dialyzed using Amicon 100 KDa cut-off centrifuge filters at least 3 times to remove residual iodixanol, after which the virus was tittered using a standard qPCR protocol^6^. AAV.FUS candidates were packaged and titered using a commercial service (Vigene biosciences) to ensure reproducibility for external investigators, as the titers can show variability between different labs^34^. We have re-titered the AAV.FUS.3 and AAV9 from another batch again in our lab, to make sure that the improvement of AAV.FUS over AAV9 is consistent between investigators.

### In vivo selection and gene delivery

To enable in vivo selection of AAV.FUS we delivered the AAV library to one hemisphere through FUS-BBBO. We targeted four sites corresponding to the striatum, dorsal hippocampus, ventral hippocampus, and midbrain using MRI guidance. We used 0.33MPa pressure and other parameters as described in the *Focused ultrasound equipment and BBB opening procedures* section. The parameters used were identical during the in vivo selection and testing of the AAV.FUS candidates. The AAVs were delivered intravenously. For the first round of selection the dose delivered was 6.7E9 viral particles per gram of body weight. The library for the first round of evolution contained 1.3E9 sequences, yielding approximately 5 particles of each clone per gram of body weight. For the second round, where the library contained 2098 candidates, 1.3E9 viral particles per gram of body weight were delivered, yielding 6.2E5 viral particles for each clone per gram of body weight. After FUS-BBBO mice were returned to the home cages for 14 days, after which they were euthanized by CO^2^ overdose.

### Tissue preparation for DNA extraction

The brains of mice euthanized with CO^2^ overdose were extracted, and the targeted hemisphere was separated from the control hemisphere with a clean blade. Each hemisphere was then frozen at −20C prior to DNA extraction. The brains were then homogenized in Trizol using a BeadBug tissue homogenization device with dedicated pre-filled 2.0 ml tubes with beads (Zirconium coated, 1.5 mm, Benchmark Scientific, Sayreville, New Jersey) for 1-3 minutes until tissue solution was homogenous. The DNA was then extracted with Trizol and amplified first with CRE-independent, and then CRE-dependent PCR as previously published^6^.

### Next Generation Sequencing Data Analysis

Next generation sequencing was performed using MiSeq (Illumina, San Diego, CA) using paired-end 75-base pair reads. The variable region of all detected capsid sequences was extracted from raw fastq files using the awk tool in Unix terminal. This process filtered out sequences not containing the constant 19bp region flanking each side of the variable region. Sequences were then sorted, checked for length, and ordered from highest to lowest copy number in the sequencing experiment. During the first screen, the top 3000 were chosen. Among these 3000, any sequence that was only a point mutation away from a sequence and 30x less abundant was removed and assumed to be a potential sequencing readout error. This led to our final library of 2098 sequences, which were synthesized by Twist Biosciences (San Francisco, CA) for use in the second round of screening. This second AAV library also included a set of 2098 “codon-optimized” capsid variants that were encoded for the same protein as the original sequences but using a different DNA sequence chosen by the IDT codon optimization tool. To process the second batch of sequencing data, we first normalized the copy numbers of the sequences in each experiment to one to ensure comparability of different samples. Then, we filtered out sequences that were not contained within the input library. Finally, we evaluated the normalized frequency of reads for each sequence, defined as the normalized copy number of each sequence averaged among original and codon-optimized variants for each capsid. Top sequences for further analysis were selected to be the most abundant sequences that appeared at least 100x more frequently in the targeted brain hemisphere than the non-targeted hemisphere in all tested mice, and from these sequences, the top 5 were chosen as AAV.FUS candidates.

### Histology, and image processing

After cardiac perfusion and extraction brains were post-fixed for 24 h in neutral buffered formalin (NBF). Brains were then sectioned at 50-micron on Compresstome VF-300 (Precisionary Instruments, Natick, MA). Sections were immunostained with an anti-NeuN Alexa Fluor 405-conjugated antibody (RBFOX3/NeuN Antibody by Novus Biological, Littleton, CO, USA, stock number: NBP1-92693AF405). Sections were imaged on a Zeiss LSM-800 microscope using a 20x objective. Channels’ laser intensities normalized to the brightness of mCherry and GFP proteins. Images were then randomized, anonymized, and exported as greyscale to ensure a lack of bias in color perception. The experimenter was blinded in terms of fluorophore color, tested AAV strain, or the mouse identification (H.L.). Three 100-micron sections of the brain were analyzed for each mouse, for each strain of the AAV including the section at the center of the FUS-target and the sections 500 and 1000 microns anterior to that section. The data was then independently validated by an experimenter blinded to the goals of the study (J.T). The inter-experimenter variability was 12.5% (1.9-fold (RL, primary scorer) vs 2.1-fold difference (JT, secondary scorer), n=15 randomly selected images, a total of 11,230 cells counted) and the difference between the scores was not statistically significant (p=0.071, two-tailed, paired t-test). To evaluate the BBB permeability of the AAV in the absence of FUS-BBBO (off-target transduction), a randomly chosen untargeted region at least 2 mm from the center of targeted region (4 times the distance of distance half-width half maximum of pressure, resulting in ^~^16-fold pressure reduction) was used within the same sections that were used to evaluate transduction efficiency at FUS focus.

### Statistical analysis

Two tailed t-test, without assuming equal variance, was used when comparing means of two data sets. For comparison of more than two data sets, one-way ANOVA was used, with a Tukey’s HSD post-hoc test to determine significance of pairwise comparisons. When more than one variable was compared across multiple samples, two-way ANOVA was used, followed by Sidak’s multiple comparisons test.

## ACKOWLEDGEMENTS

The authors thank Drs. Benjamin Deverman, Nicholas Flytzanis, Nicholas Goeden, and Viviana Gradinaru, and the CLOVER center at Caltech for helpful discussions. This research was supported by the National Institutes of Health (grant UG3MH120102 to MGS), the Jacobs Institute for Molecular Engineering in Medicine and the Sontag Foundation, and 2019 NARSAD Young Investigator Grant from the Brain and Behavior Research Foundation (grant 27737 to JOS). Related work in the Shapiro Lab is supported by the David and Lucille Packard Foundation and the Heritage Medical Research Institute and in Szablowski lab by The G. Harold and Leila Y. Mathers Charitable Foundation. JEH acknowledges support from Rose Hills foundation and Barry Goldwater Scholarship.

