## Supplementary information for "Engineering Viral Vectors for Acoustically Targeted Gene Delivery"

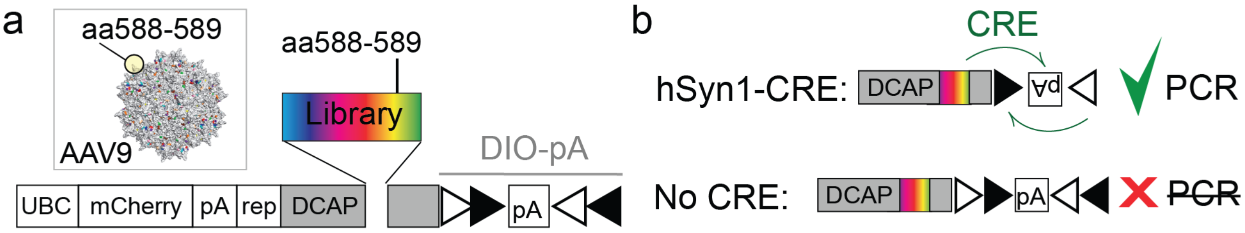


**Supplementary Figure S1. Construction of the AAV library and CRE-dependent PCR. a**, Randomized 21-basepair DNA fragment was inserted into the AAV9 capsid between aminoacids 588 and 589, which resides at the exterior of an AAV capsid (inset). AAV capsid was produced within the AAV genome allowing for recovery of the capsid sequenced from transduced cells. The capsid coding sequence was followed by a polyA (pA) sequence flanked by a double-inverted floxed open reading frame (DIO). **b**, The DIO sequence can be recombined and inverted in the presence of Cre enzyme. That sequence inversion can then be detected using PCR. Therefore, the DNA from AAVs that transduced cells expressing Cre can be amplified using a PCR reaction. In our study, we used hSyn1-Cre mice which express Cre selectively in neurons and thus we selecting for neuron-transducing AAVs.

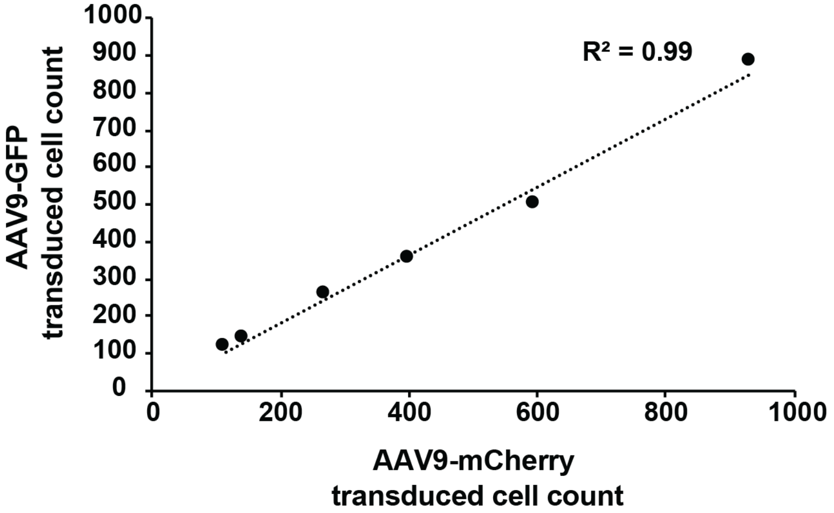


**Supplementary Figure S2. AAV9 GFP + AAV9 mCherry.** Transduced cell counts comparing AAV9 carrying GFP and mCherry are highly correlated and (Rsq=0.99). The mean numbers of transduced cells are not significantly different (fold difference between AAV9-GFP and AAV9-mCherry: 1.07-fold, p=0.081 (ns), paired t-test, 6 sections tested from 2 mice).

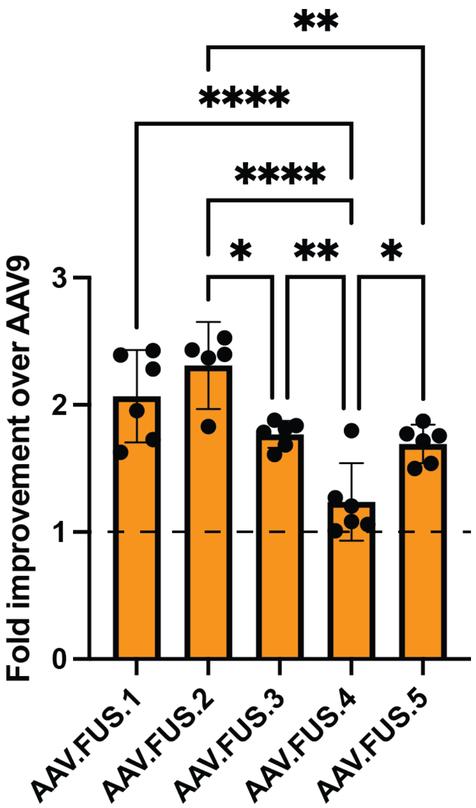


**Supplementary Figure S3. Pairwise comparison of AAV.FUS candidates transduction of the brain.** Non-significant comparisons not shown for clarity**.** (**** = p<0.0001; *** = p<0.001; ** = p <0.01; * = p<0.05; F(4, 24) = 14.96, P<0.0001, One-way ANOVA with Tukey HSD post hoc test.). Non-significant pairwise comparisons not shown for clarity. Error bars are 95% CI.

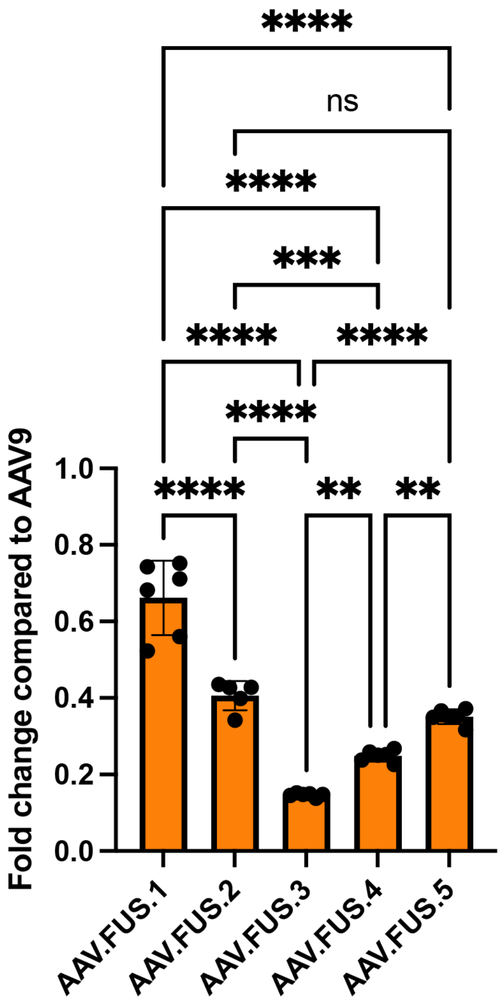


**Supplementary Figure S4. Pairwise comparison of AAV.FUS candidates transduction of the liver. Non-significant comparisons not shown for clarity.** AAV.FUS.3 shows significantly reduced liver transduction compared to other AAV.FUS candidates. One-way ANOVA with Tukey HSD post-hoc test. $F(4, 24) = 96.69$. $P<0.0001$; All pairwise comparisons are below $p<0.0001$, except AAV.FUS.2 vs AAV.FUS.4 ($p=0.0001$), AAV.FUS.2 vs AAV.FUS.5 ($p=0.3524$), AAV.FUS.3 vs AAV.FUS.4 ($p=0.01$), and AAV.FUS.4 vs AAV.FUS.5 ($p=0.0099$)). (**** = $p<0.0001$; *** = $p<0.001$; ** = $p <0.01$; * = $p<0.05$, ns = non significant). Error bars are 95% CI.

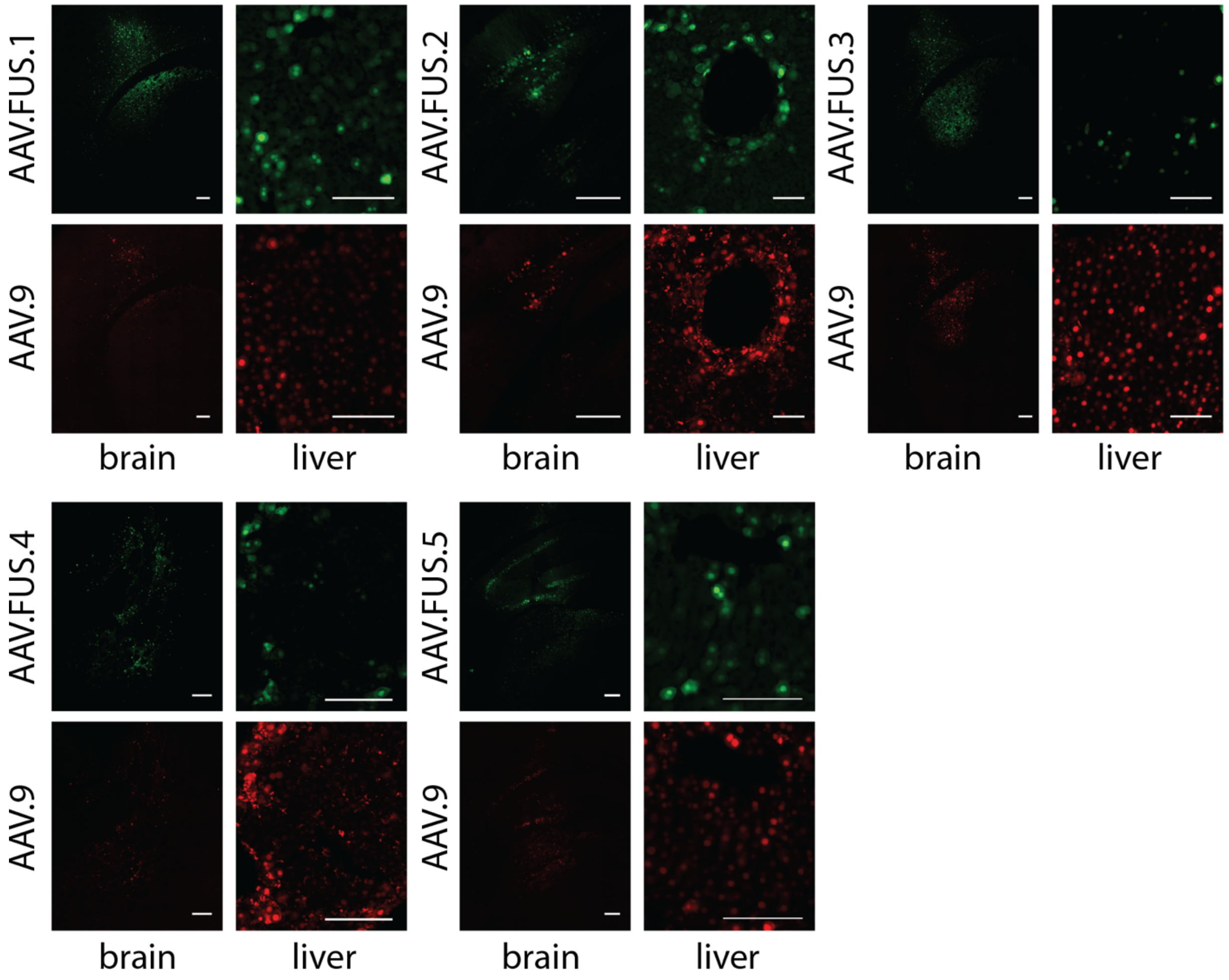


**Supplementary Figure S5. Representative images of transduction in brain and liver for all AAV.FUS (green, EGFP) and corresponding co-injected AAV9 control (red, mCherry) vectors.** Scale bars, 200 microns for the brain, 100 microns for the liver.

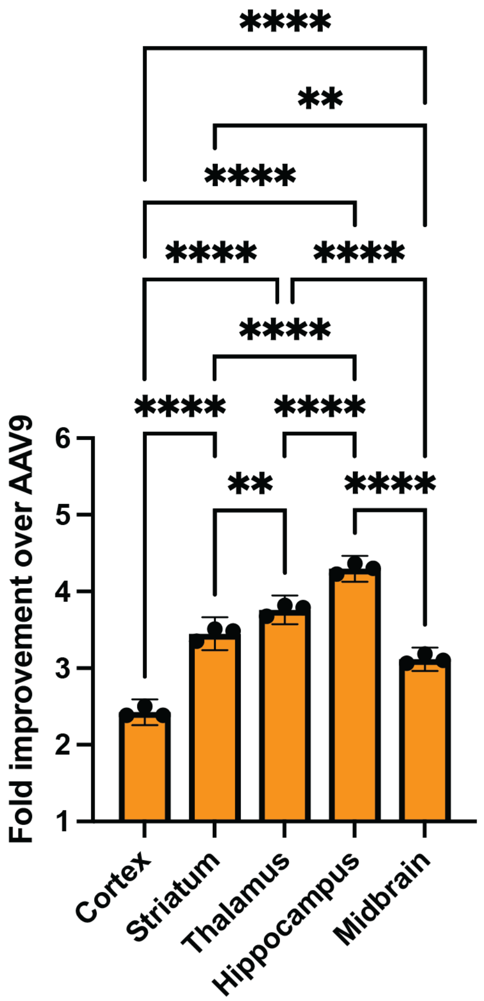


**Supplementary Figure S6. Detailed pairwise comparisons for analysis of regional dependence of transduction efficiency for AAV.FUS.3.** (**** = p<0.0001; *** = p<0.001; ** = p <0.01; * = p<0.05, ns = non significant; One-way ANOVA with Tukey HSD post-hoc test.).
